# Switch independent task representations in frontal and parietal cortex

**DOI:** 10.1101/138230

**Authors:** Lasse S. Loose, David Wisniewski, Marco Rusconi, Thomas Goschke, John-Dylan Haynes

## Abstract

Alternating between two tasks is effortful and impairs performance. Previous functional magnetic resonance imaging (fMRI) studies have found increased activity in fronto-parietal cortex when task switching is required. One possibility is that the additional control demands for switch trials are met by strengthening task representations in the human brain. Alternatively, on switch trials the residual representation of the previous task might impede the buildup of a neural task representation. This would predict weaker task representations on switch trials, thus also explaining the performance costs. To test this, participants were cued to perform one of two similar tasks, with the task being repeated or switched between successive trials. MVPA was used to test which regions encode the tasks and whether this encoding differs between switch and repeat trials. As expected, we found information about task representations in frontal and parietal cortex, but there was no difference in the decoding accuracy of task-related information between switch and repeat trials. Using cross-classification we found that the fronto-parietal cortex encodes tasks using a similar spatial pattern in switch and repeat trials. Thus, task representations in frontal and parietal cortex are largely switch-independent. We found no evidence that neural information about task representations in these regions can explain behavioral costs usually associated with task switching.

**Significance statement:** Alternating between two tasks is effortful and slows down performance. One possible explanation is that the representations in the human brain need time to build up and are thus weaker on switch trials, explaining performance costs. Alternatively, task representations might even be enhanced in order to overcome the previous task. Here we used a combination of fMRI and a brain classifier to test whether the additional control demands under switching conditions lead to an increased or decreased strength of task representations in fronto-parietal brain regions. We found that task representations are not significantly modulated by switching processes. Thus, task representations in the human brain cannot account for the performance costs associated with alternating between tasks.

## Introduction

In order to reach desired goals, humans are often required to switch between different tasks. This important aspect of cognitive control allows flexible adjustment of behavior to changing circumstances (Kok, Ridderinkhof, & Ullsperger, 2006). Such adjustments are often investigated using the task switching paradigm, requiring subjects to frequently switch between two or more tasks (Meiran, 2010). Typically, participants react slower and perform less accurate on tasks they just switched to as compared to tasks that were repeated multiple times (Jersild, 1927; Spector & Biederman, 1976). These *switch costs* (Rogers & Monsell, 1995) reflect cognitive control processes (Goschke, 2000) that affect task processing and the implementation of tasks (Monsell, 2003), as well as proactive interference and between-task crosstalk (Allport, Styles, & Hsieh, 1994; Yeung, 2006). However, the exact sources of switch costs are still under debate (Kiesel et al., 2010).

Previous fMRI studies investigated the neural basis of preparatory processes in task switching using univariate methods (Ruge, Jamadar, Zimmermann, & Karayanidis, 2011). Although many results implicate involvement of prefrontal and parietal cortical regions in task switching (Dove, Pollmann, Schubert, Wiggins, & Yves von Cramon, 2000; Gruber, Karch, Schlueter, Falkai, & Goschke, 2006; Jamadar, Hughes, Fulham, Michie, & Karayanidis, 2010), this finding is not always consistent (Ruge et al., 2011). Previous task switching research mostly focused on neural correlates of task switching processes in terms of BOLD signal differences between switch and repeat trials. Recently, multivoxel pattern analysis (MVPA, Haynes, 2015) has been used to investigate neural task representations. Such representations are encoded in local spatial activation patterns in the lateral prefrontal, dorsal anterior cingulate and posterior parietal cortex (Bode & Haynes, 2009; Gilbert, 2011; Wisniewski, Reverberi, Momennejad, Kahnt, & Haynes, 2015; Woolgar, Hampshire, Thompson, & Duncan, 2011).

Different cognitive processes like rule complexity (Woolgar, Afshar, Williams, & Rich, 2015) or skill acquisition (Jimura, Cazalis, Stover, & Poldrack, 2014) have been shown to alter representations of tasks. However, whether and how task switching (and its associated cognitive control demands) influence task representations is still largely unknown. Behavioral switch costs in task switching reflect both the cognitive control processes required to switch to performing a different task as well as involuntary processes such as proactive interference from a previous task-set (Kiesel et al., 2010; Vandierendonck, Liefooghe, & Verbruggen, 2010). Possibly, this also affects the representation of these tasks (Waskom, Kumaran, Gordon, Rissman, & Wagner, 2014). In other cases, task representations remain unaffected by whether tasks were chosen freely or were externally cued (Wisniewski, Goschke, & Haynes, 2016) or whether tasks were novel or routinely performed (Cole, Etzel, Zacks, Schneider, & Braver, 2011). This suggests that tasks can also be represented independently of current cognitive control demands (Zhang, Kriegeskorte, Carlin, & Rowe, 2013). However, it has remained open whether and how task-switch related control demands and between-task crosstalk in task-switching contexts affect the neural representation of tasks.

In order to investigate the influence of task switching on task representations two main questions are addressed in this study: (1) Do different cognitive control demands on task-switch versus task-repeat trials affect the strength of neural tasks representations? More specifically, does the accuracy with which tasks can be decoded from neural activation patterns differ between task-switch and task-repeat trials? (2) Is the neural code in which tasks are represented independent from control demands? Thus, are tasks encoded using similar spatial activation patterns in switch and repeat trials?

To address these questions, subjects were instructed to perform one of two simple stimulus-response mapping tasks while brain activity was measured with fMRI. We identified brain networks involved in representing tasks and then assessed task information in these regions for switch vs. repeat trials separately. Furthermore, we tested whether brain regions encode tasks similarly in switch and repeat trials. Results indicated that tasks are represented similarly in a fronto-parietal network, suggesting that switch-related cognitive control demands exert no strong effect on neural task representations.

## Materials and Methods

### Participants

42 right-handed subjects (21 females, mean age: 25.2, age range 20-30 years) with normal or corrected to normal vision participated in the study. We obtained written informed consent from each subject and the local ethics committee approved the experiment. Subjects received 30€ for their participation. No subject had a self-reported history of neurological or psychiatric disorders. We only invited subjects to the fMRI session whose accuracy in performing the tasks after training was above 90 % and we thus had to discard one subject after the training session because of poor behavioral performance (see experimental paradigm). We discarded two further subjects because of technical problems during scanning and one subject due to excessive head movement during scanning. To ensure reliable behavioral performance all subjects took part in a training session 1-3 days prior to the scanning. Overall, the MRI data of 38 subjects (20 females, mean age: 25, age range 20-29 years) were used for our analyses.

### Task and Experimental Paradigm

Subjects were cued to apply one of two stimulus-response mappings (tasks) to a visual stimulus in each trial of the experiment. In half of the trials, the task was identical to the previous trial (repeat trials), in the other half of the trials the task differed from the previous trial (switch trials). We instructed subjects to respond as quickly and accurately as possible.

The experiment was programmed using MATLAB Version 7.11.0 (MathWorks, RRID: SCR_001622) and the Cogent Toolbox (http://www.vislab.ucl.ac.uk/cogent.php). On each trial we first presented a task screen for 3,000 ms that simultaneously displayed a task cue, a target stimulus and four colored circles used for response mapping assignment (Figure 1 and see below). Subjects were allowed to respond in the same 3,000 ms time window. The task screen was followed by an intertrial-interval (ITI) where a fixation cross was presented centrally on screen. ITIs varied between 4,000, 6,000, 8,000 and 10,000 ms and were distributed pseudo-logarithmically to decorrelate trials in time. The mean ITI was 5,525 ms.

**Figure 1.**
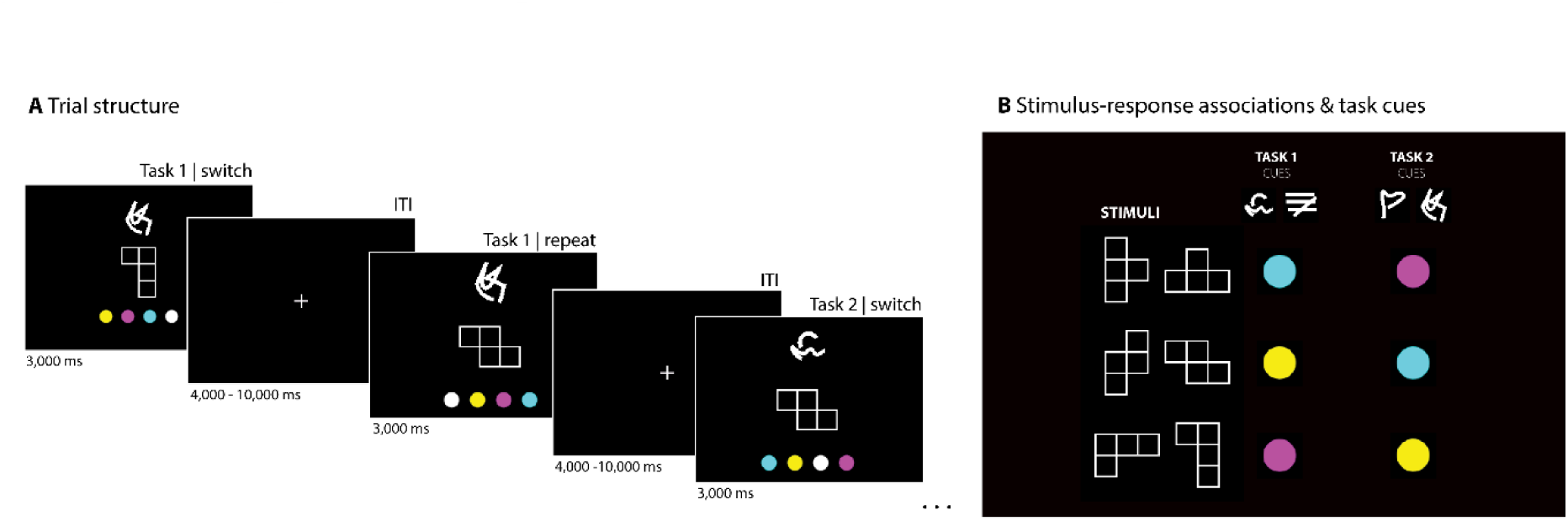
Experimental paradigm. **A** Trial structure. Each trial consisted of a target stimulus at fixation, a cue stimulus above, and four possible response options below. Each task screen was presented for 3,000 ms, during which participants could indicate a response using the left and right index and middle fingers. Each trial was followed by a fixation cross with a variable inter-trial-interval (ITI, 4,000-10,000 ms, mean = 5,525 ms). Responses were indicated by pressing the button corresponding to the mapped color on screen on a 2 × 2 button box with their index and middle fingers of both hands. Subjects were cued to perform one of two tasks, switching between tasks or repeating a task up to three consecutive times. **B** Stimulus-response associations & task cues. The two tasks consisted of similar stimulus response mappings, associating stimulus shapes (in two possible orientations) with colors. Each task was indicated by one of two possible abstract cues.

Tasks were cued using abstract visual symbols presented at the top of the screen. They were free of semantic meaning in order to avoid a-priori semantic associations (Figure 1, Reverberi, Görgen, & Haynes, 2012a; Wisniewski, Reverberi, Momennejad, et al., 2015). Over the experiment, two different cues were associated with each task in order to allow for cue-independent task decoding (see below for details, also see Reverberi, Görgen, & Haynes, 2012). The cue - task associations were counterbalanced across subjects. The target stimuli consisted of three geometric objects (Figure 1, T-shape, L-shape, Z-shape) each appearing in two possible orientations (0 and 90 degrees) and presented in the middle of the screen. Stimuli and their orientations were pseudo-randomized in order to control for the influence of low-level visual features. The two tasks consisted of different stimulus response (S-R) mappings that associated stimulus shapes with colors that in turn specified which response key had to be pressed. In task one, the T-shaped stimulus was associated with magenta, the Z-shaped stimulus with cyan and the L-shaped stimulus with yellow. In task two, the T-shaped stimulus was associated with cyan, the Z-shaped stimulus with yellow and the L-shaped stimulus with magenta. The S-R mappings were chosen to be similar in order to control for possible confounds due to difficulty differences between tasks (Todd, Nystrom, & Cohen, 2013). Furthermore, switch costs can also be affected by task difficulty (Arbuthnott, 2008). Below the target stimulus four colored circles were presented that mapped colors to response buttons. The position of the each colored circle was pseudo-randomized across trials, avoiding motor preparation of responses as well as balancing left-hand and right-hand button presses throughout the experiment. Subjects used index and middle fingers of both hands to indicate their response by pressing the button corresponding to the color on screen on a 2 × 2 button box (Current Designs). Three of the circles were relevant for the task (cyan, magenta, yellow) and one was a dummy (white) that was not relevant in any trial. This was done in order to balance left and right button presses.

Each run contained 80 trials, which were ordered so that 50 % appeared in a sequence-length of 1 (e.g. task1), 37.5 % in a sequence length of 2 (e.g. task 1 – task 1), and 12.5 % in a sequence length of 3 (e.g. task 1 – task 1 – task 1). This results in 50 % switch trials and 50 % repeat trials overall. In 50 % of the trials subjects performed task 1 and in 50 % they performed task 2. Tasks and switch conditions were orthogonalized. Within each brief sequence of identical tasks we only used one of the two possible cues, i.e. in each of the subsequent repetitions of a task the same cue was used (cue repetition). Furthermore, cues were counterbalanced with stimuli and ITIs, to avoid possible confounds.

Following each completed run, the percentage of correct answered trials was presented and subjects were offered a short resting break of self-determined length. Subjects performed 5 runs in total. The experiment lasted around 75 minutes in total. A sixth run, in which subjects performed the tasks in a different sequential order, was not analyzed and is not included in this paper.

1-3 days before the scanning session, subjects performed a 90 minutes training session, during which they learned the S-R mappings. At the end of the training session, they performed two runs of the task as they would be presented in the scanner. Only if the accuracy during these runs was above 90 % were subjects invited to the scanning session. This was done in order to avoid possible learning effects during the MRI session.

### Image Acquisition

Functional imaging was conducted on a 3-T Siemens Trio (Erlangen, Germany) scanner equipped with a 12-channel head coil. For each of the 5 relevant scanning sessions we acquired 347 T2*-weighted (TR = 2000 ms; TE, 30 ms; flip angle, 90°) gradient-echo echo-planar images (EPI). Imaging parameters were as follows: repetition time (TR), 2000 ms; echo time (TE), 30 ms; flip angle, 90°. Each volume contains 33 slices (thickness: 3 mm) separated by gaps of 0.75 mm. Matrix size was 64 × 64, the field of view (FOV) was 192 mm and in-plane voxel resolution was set to 3 mm² with a voxel size of 3 × 3 × 3 mm. A T1-weighted structural dataset was also collected. The parameters were as follows: TR, 1900 ms; TE, 2.52 ms; matrix size, 256 × 256; FOV, 256 mm; 192 slices (1 mmt thick); flip angle, 9°.

### Data Analysis

In all analyses only trials with correct responses and preceded by correct trials (no misses/errors) were included in order to avoid post-error slowing effects (Dudschig & Jentzsch, 2009). We analyzed behavioral and fMRI-data using MATLAB Version 2013a (MathWorks), and for the multivariate analyses we used *The Decoding Toolbox* (TDT, Hebart, Görgen, Haynes, & Dubois, 2015). Unthresholded group-level parametric maps of all analyses can be found at NeuroVault (Gorgolewski et al., 2015, RRID:SCR_003806; http://neurovault.org/collections/2011/).

### Behavior

For each subject we assessed task performance by calculating the mean RT and mean accuracy (percentage of trials that were correctly answered in time) across all runs. It has been reported previously, that switching between tasks leads to increased RT and decreased accuracy in switch trials compared to repeat trials (Monsell, 2003). We tested these so-called switch cost*s* by comparing switch and repeat trials in terms of mean RT and accuracy. In order to control for possible influences of task difficulty we also assessed the influence of the two tasks and the four cues on RTs and accuracies. We expected task switches to have an effect on both accuracy and RT (switch cost) but did not expect the other variables to affect them.

### Neuroimaging

#### First level GLM analysis

In a first step, we analyzed functional data using SPM8 (http://www.fil.ion.ucl.ac.uk/spm, RRID: SCR_007037). The functional volumes were unwarped, realigned and slice time corrected. No spatial smoothing and no spatial normalization was applied at this point in order to preserve fine-grained patterns of voxel activations (Haynes & Rees, 2006).

The preprocessed data was used to estimate a voxelwise general linear model (GLM; Friston, Jezzard, & Turner, 1994). Twelve regressors-of-interest were used in the GLM. First, regressors for the 8 conditions of the experimental design: 2 (tasks) × 2 (cue-sets) × 2 (switch / repeat) were added. Second, 4 separate regressors of no interest were added modelling the four possible button presses in order to control for possible motor confounds in the data. Third, movement parameters were added to the GLM as regressors of no interest in order to account for possible head movement during scanning. Regressors were time-locked to the onset of the task-screen and convolved with a canonical hemodynamic response function (HRF), as implemented in SPM.

In order to account for the possible influence of task difficulty on MVPA results (Todd et al., 2013) we first calculated the mean RT for task 1 and task 2 for each subjects individually. We then set the duration of each regressor to the mean task RT of the current trial (mean RT task 1 for trials with task 1, and mean RT task 2 for trials with task 2, as suggested by Woolgar, Golland, & Bode, 2014). This accounts for task specific RT related effects in the data during GLM estimation but does not remove task switch related variance from the data (for recent reviews about switch cost see Kiesel et al., 2010; Vandierendonck et al., 2010).

#### Multivariate searchlight decoding

##### Analysis 1: Differences in task coding in switch and repeat trials

In order to test for possible differences of task representations in switch and repeat trials we first identified regions that code for tasks and, in the following steps, assessed the differences of task-decoding in switch and repeat trials separately in these regions.

**Analysis 1A. Task information across all trials:** In the first analysis, we used “searchlight” MVPA (Kriegeskorte, Goebel, & Bandettini, 2006; Norman, Polyn, Detre, & Haxby, 2006) as implemented in TDT (Hebart et al., 2015) on the maps of GLM-parameter estimates for each individual subject. For each voxel V in the volume the searchlight classifier distinguishes between the two classes (here: tasks) based on the multivariate pattern formed by the local fMRI activity patterns in a small spherical cluster with the radius of 3 voxels surrounding V. We used a support vector classifier (SVC) with a linear kernel and a fixed regularization parameter (C = 1) as implemented in LIBSVM (http://www.csie.ntu.edu.tw/∼cjlin/libsvm). As a result, searchlight decoding produces a whole-brain accuracy map, representing which searchlights contained information about the two classes entered into the analysis. To identify which brain regions contain information about tasks, we performed this first searchlight decoding analysis, classifying task 1 vs task 2 and using data from both switch and repeat trials combined. Trials were collapsed across switch and repeat condition in order to increase power to identify regions of interests (ROI) that contain information about tasks. In order to control for the effect of visual cue information, we performed cross-classification across visual cues. More specifically, we trained the SVC to discriminate “task 1 with cue 1” and “task 2 with cue 2”, and tested its performance on trials from “task 1 with cue 3” and “task 2 with cue 4”. Only brain regions that use similar activation patterns to encode the same tasks with different cues will be visible in this analysis. Therefore, this analysis controls for effects that are merely due to the visual features of the cues used. There are a total of four different combinations of task and visual cues as training- and test-dataset, so that we repeated this analysis three more times: once for every combination. In order to address the problem of overfitting (Kriegeskorte, Simmons, Bellgowan, & Baker, 2009), we performed a five-fold leave-one-run-out cross-validation (CV). Thus, every run was the test dataset once. The results of the combinations of cross-validation and cross-classification were averaged for each subject.

The average accuracy maps were then spatially normalized to a standard brain (Montreal Neurological Institute [MNI] EPI template of SPM8) to account for individual differences in brain structure. Accuracy maps were then smoothed with a Gaussian kernel (6mm full-width at half-maximum) in order to account for differences in localization. At the group level, a random-effects analysis was performed, using voxel-wise one sample t-tests against chance level (50 %). Results were initially thresholded at voxel level with *p* < 0.001, corrected for multiple comparisons at the cluster level for familywise error (FWE, *p* < 0.05). Note, that these threshold values are not problematic for cluster-level inference regarding the inflated FWE-rates that have recently been discovered by Eklund, Nichols, & Knutsson, 2016.

**Analysis 1B. Differences in task decoding for switch and repeat trials:** In a second step, we performed two additional searchlight decoding analyses that were highly similar to analysis 1A described above. This time we performed two independent analyses for switch trials only and repeat trials only. We first entered only the data of switch trials into a SVC that was trained to classify task 1 versus task 2. We again applied cross-classification across cues and leave-one-run-out cross-validation and averaged across them. We also smoothed and normalized the resulting decoding accuracy maps, as described above. The same procedure was repeated for repeat trials only. This yielded a task decoding accuracy map for switch trials and for repeat trials for each individual subject. To compare the task decoding accuracies in switch and in repeat trials we created regions of interest (ROIs) from the clusters that we defined in task decoding analysis 1A. In order to avoid circular analysis (Kriegeskorte et al., 2009), we used a leave-one-subject-out ROI analysis (Esterman, Tamber-Rosenau, Chiu, & Yantis, 2010). For this, we excluded one subject and performed a group level analysis as described above (analysis 1A). The results were then thresholded at voxel level with *p* < 0.001 (corrected for multiple comparisons at the cluster level, FWE, *p* < 0.05). We extracted the resulting significant clusters from this analysis and created a ROI from each cluster (based only on the training subjects). For each ROI thus defined, we extracted the mean decoding accuracy for the left out subject. The ROI should resemble the group level results of analysis 1A, but ensure an independent dataset for extracting decoding accuracies. Accuracy values were extracted for the decoding of task in switch trials only, repeat trials only, and all trials together (analysis 1A). We repeated this procedure until every subject was left out once. This ensures independence of the data used to define the ROIs from the data used for statistical assessment of the accuracy values inside these ROIs. The mean decoding accuracies from all three analyses and all ROIs were then entered into a two-factorial repeated measures ANOVA (Factor 1: 3 analyses, Factor 2: ROIs) in order to identify possible differences between task coding in switch and repeat trials in each ROI. Furthermore, in order to assess whether decoding accuracies were significantly above chance in each analysis and ROI, planned one-tailed t-test against chance level were performed. Results from these tests were Bonferroni corrected for the three analyses performed in each ROI.

##### Analysis 2: Similarities in task coding between switch and repeat trials

Please note that the abovementioned analysis (1D) merely tests whether brain regions that encode tasks have different accuracies in switch and in repeat trials. If a given ROI indeed has a higher accuracy in one or the other condition, this would indicate a *specialized* role for task coding in either switch or repeat trials. If however, no difference were to be found, this would not directly show that the ROI has a *similar* role in switch and repeat trials. In order to directly assess whether any brain regions encode tasks similarly in these two conditions, a different type of analysis is necessary. Thus, in analysis 2, we aimed to identify brain regions that encode task-information in the same way independent of whether subjects were repeating or switching between tasks, again using cross-classification (Kaplan, Man, & Greening, 2015; Reverberi et al., 2012a; Wisniewski et al., 2016). Similar to analysis 1A, we first trained a searchlight classifier to distinguish between tasks in switch-trials only and tested it on repeat trials only. We then trained a classifier on repeat trials only and tested it on switch trials only. Again, in both cases we used leave-one-run-out cross-validation in order to avoid the problem of overfitting. Results from both analyses were first averaged for both cross-classification directions and then smoothed and normalized as in the previous analyses.

Please note that in contrast to the analysis 1, this analysis does not control for the effect of visual features of the task cues, and results might potentially reflect these. Due to the limited number of trials, we were unable to perform a cross-classification across switch/repeat and visual cues at the same time. In order to still control for the effect of visual cue information we again used the ROIs defined in analysis 1A, using the leave-one-subject-out method. Within these clusters we now extracted the mean task decoding accuracies from analysis 2, where we cross-classified across switch and repeat trials. Please note, that this is similar to a conjunction analysis, and only voxels that show significant above chance information in both task decoding cross-classified across visual cues and task decoding cross-classified across the switch and repeat conditions are interpreted. If tasks are encoded similarly in these regions, mean decoding accuracies of task in both analyses should be significantly above chance. We tested this by applying a t-test (against chance level, 50 %) on the mean decoding accuracies for each cluster.

## Results

### Behavior

The mean RT across all correct trials was 1,681 ms (SE = 30 ms). After removing trials following error trials the mean RT changed significantly to 1,664 ms (SE = 27 ms; paired t-test: *t*(37) = 3.69; *p* < 0.001). This effect could reflect post error slowing (Dudschig & Jentzsch, 2009). All fMRI and RT analyses are based only on correct trials also following a correct trial. On average, subjects were correct and fast enough in 95.5 % (SE = 0.6 %) of the trials. In 2.9 % of the trials (SE = 0.3 %) subjects pressed the wrong button and in the 1.6 % (SE = 0.3 %) they did not respond within the 3,000 ms response window. The mean RT did not differ significantly between the two tasks (paired t-test, *t*(37) = −0.30, *p* = 0.76), neither did the accuracy of both tasks. Furthermore, there was no significant effect of cue on RTs, as tested using a one-way repeated measures (ANOVA, *F*(3, 37) = 0.31, *p* = 0.81). No effects of tasks (paired t-test, *t*(37) = 0.47, *p* = 0.74) or cues (ANOVA, *F*(3, 37) = 1.17, *p* = 0.32) were found in accuracy rates. The average RT in switch trials was 1,699 ms (SE = 32 ms). The average RT in repetition trials was 1,656 ms (SE = 30 ms). The difference between these switch and repeat trials (switch cost) was significant, (paired t-test, *t*(37) = 5.04; *p* < 0.001). Average accuracy in switch trials was 94.59 % (SE = 0.68 %) and in repeat trials 96.36 % (SE = 0.5 %). This difference was also significant (paired t-test, *t*(37) = −4.44; *p* < 0.001). These results replicate previous findings of switch cost in RT and accuracy values (Monsell, 2003).

#### Multivariate searchlight decoding

##### Analysis 1: Differences in task coding in switch and repeat trials

**Analysis 1A. Task information across all trials:** First, we identified regions which encode tasks using data from both switch and repeat trials combined. Using cross-classification, we ensured that the visual features of the task cues cannot explain the results. Significant above-chance classification of task could be observed in three clusters (*p* < 0.05, FWE corrected at the cluster level, initial voxel threshold *p* < 0.001, Figure 2A and Table 1): the first is located in left inferior and superior parietal cortex spanning across angular gyrus; the second cluster was found in right superior parietal cortex spanning across angular gyrus; the third cluster is located in left prefrontal cortex (PFC).

**Figure 2.**
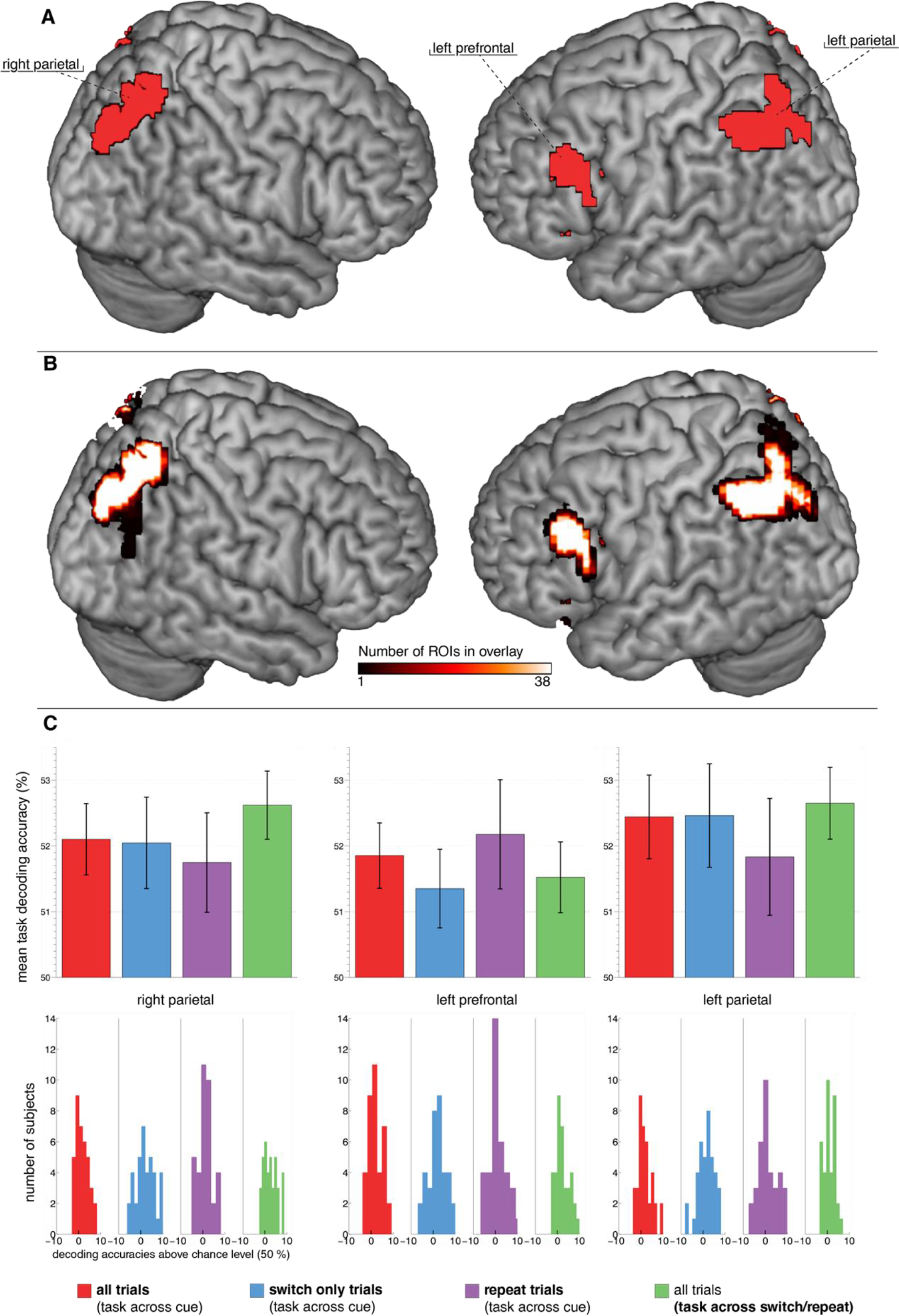
Task decoding. **A** Task decoding across cues in all trials and all subjects. Tasks were encoded in bilateral superior parietal cortex, left inferior parietal cortex and left lateral prefrontal cortex (*p* < 0.05, FWE corrected at the cluster level, initial voxel threshold *p* < 0.001). **B** Overlay of all 38 leave-one-subject-out ROIS. All ROIs were created leaving out one subject at the group level statistic (*p* < 0.05, FWE corrected at the cluster level, initial voxel threshold *p* < 0.001) and later used for extraction of mean decoding accuracy values in that subject. **C** Mean task decoding accuracies extracted from the ROIs depicted in Figure 2B. We extracted values from four different decodings: task across cues in all trials (red), task across cues in switch trials only (blue), task across cues in repeat trials only (violet) and task across switch (green). Chance level in these plots is 50 %. The distribution of mean decoding accuracies across subjects is shown in the histograms below.

**Table 1:** Results of analysis I: Brain regions where tasks could be decoded in an analysis collapsed across switch and stay trials, independent of visual cue. Results of analysis II: Brain regions where classifiers trained on switch trials could be used to decode the task in repeat trials and vice versa.

| Brain region | Side | Cluster size | MNI coordinates (peak voxels) |  |  | t-score peak |
| --- | --- | --- | --- | --- | --- | --- |
|  |  |  | x | y | z |  |
| **I. Task across cues in all trials** |  |  |  |  |  |  |
| Parietal lobe | left | 383 | -51 | -52 | 49 | 4.75 |
| Parietal lobe | right | 293 | 36 | -61 | 64 | 4.87 |
| Prefrontal lobe | right | 261 | -39 | 35 | -2 | 5.51 |
| **II. Task across switch in all trials** |  |  |  |  |  |  |
| Parietal lobe | left | 1053 | -48 | -55 | 49 | 5.17 |
| Parietal lobe | right | 597 | 27 | -64 | 46 | 5.71 |
| Occipital lobe | left | 440 | -21 | -91 | -2 | 4.69 |
| Cerebellum | left | 173 | -39 | -64 | -23 | 5.34 |
| Occipital lobe, cerebellum | right | 135 | 45 | -64 | -11 | 4.8 |

**Analysis 1B. Differences in task decoding for switch and repeat trials:** In order to compare the task decoding accuracies in switch only and repeat only trials, we used a leave-one-subject-out approach to create the ROIs from the clusters identified in analysis 1A. This procedure avoids the problem of double-dipping (Kriegeskorte et al., 2009). We then extracted the task decoding accuracy values in switch only and repeat only conditions. Figure 2B shows an overlay of all leave-one-subject-out-ROIs that were created. As expected, they closely resemble task decoding results across all subjects in analysis 1A. A two-factorial repeated measures ANOVA on the mean task decoding accuracies in these ROIs showed no significant main effect of the decoding analysis (all/switch-only/repeat-only task decodings, *F*(2,74) = 0.06, *p* = 0.94); no significant main effect of the ROI (*F*(2,74) = 0.59, *p* = 0.55); and no interaction effect between ROI and the decoding analysis (*F*(4,148) = 1.08, *p* = 0.36). This indicates that there are no strong differences in the task decoding accuracies between switch and repeat trials in task-related brain regions.

Average task decoding in the *left parietal cortex* in all trials was 52.44 % (SE = 0.64 %), which was significantly above chance level (50 %, t-test: *t*(37) = 3.82; *p* < 0.001). In switch trials only the average decoding accuracy was 52.46 % (SE = 0.79 %) and in repeat trials only it was 51.83 % (SE = 0.89 %). In *right parietal cortex* the task decoding accuracy in all trials was 52.1 % (SE = 0.54 %), and was significantly above chance level (t-test: *t*(37) = 3.87; *p* < 0.001). In switch trials only it was 52.05 % (SE = 0.69 %) and in repeat trials only it was 51.75 % (SE = 0.76). In *left lateral prefrontal cortex* the task decoding accuracy in all trials was 51.85 % (SE = 0.5 %) and was significantly above chance level (t-test: *t*(37) = 3.733; *p* < 0.001), in switch trials only it was 51.35 % (SE = 0.6 %) and for repeat trials only 52.18 % (SE = 0.83 %).

##### Analysis 2: Similarities in task coding between switch and repeat trials

In analysis 1, we did not find evidence for a difference in task coding between switch and repeat trials. In a next step, we directly assessed whether regions that encode task do so similarly across both switch and repeat trials. We thus performed a task decoding analysis, training on switch trials and testing on repeat trials. To ensure an independent test dataset, we again used the ROIs extracted from analysis 1A using a leave-one-subject-out approach. We extracted the mean decoding accuracy in these ROIs from the task decoding analysis cross-classified across the switch/repeat conditions. Mean decoding of task was significantly above chance-level (50 %) in left parietal (t-test: *t*(37) = 4.84; *p* < 0.001), right parietal (t-test: *t*(37) = 5.05; *p* < 0.001) and left prefrontal (t-test: *t*(37) = 2.83; *p* < 0.001)) regions. This finding indicates that all identified task-related brain regions encode tasks similarly regardless of the current switch/repeat condition.

In order to assess whether any other regions outside of the ROIs investigated above also encode tasks similarly in switch and repeat trials, we performed an additional explorative whole-brain analysis of the task decoding using cross classification across switch / repeat trials.

Results were thresholded at voxel level with *p* < 0.001, corrected for multiple comparisons at the cluster level (FWE, *p* < 0.05). Task information was found in bilateral inferior and superior parietal cortex, bilateral precuneus, right angular gyrus and bilateral occipital cortex spanning into bilateral cerebellum (Figure 3, green). Please note that in contrast to analysis 1, this analysis does not control for the effect of visual features of the task cues, and results might potentially reflect these. We therefore performed a conjunction analysis with the regions identified in analysis 1A. This analysis explicitly controls for the influence of visual cue features on task decoding results. Voxels found in both analysis 1A *and* analysis 2 thus encode tasks similarly for different visual cues and different switch/repeat conditions. This conjunction analysis identified the bilateral parietal cortex (Figure 3, yellow). In contrast to analysis 1A, we did not find a prefrontal cluster. Please note that this whole-brain analysis is less sensitive than our leave-one-subject-out ROI approach, potentially explaining the absence of prefrontal findings. This analysis suggests that the parietal cortex encodes tasks similarly across multiple different contexts.

**Figure 3.**
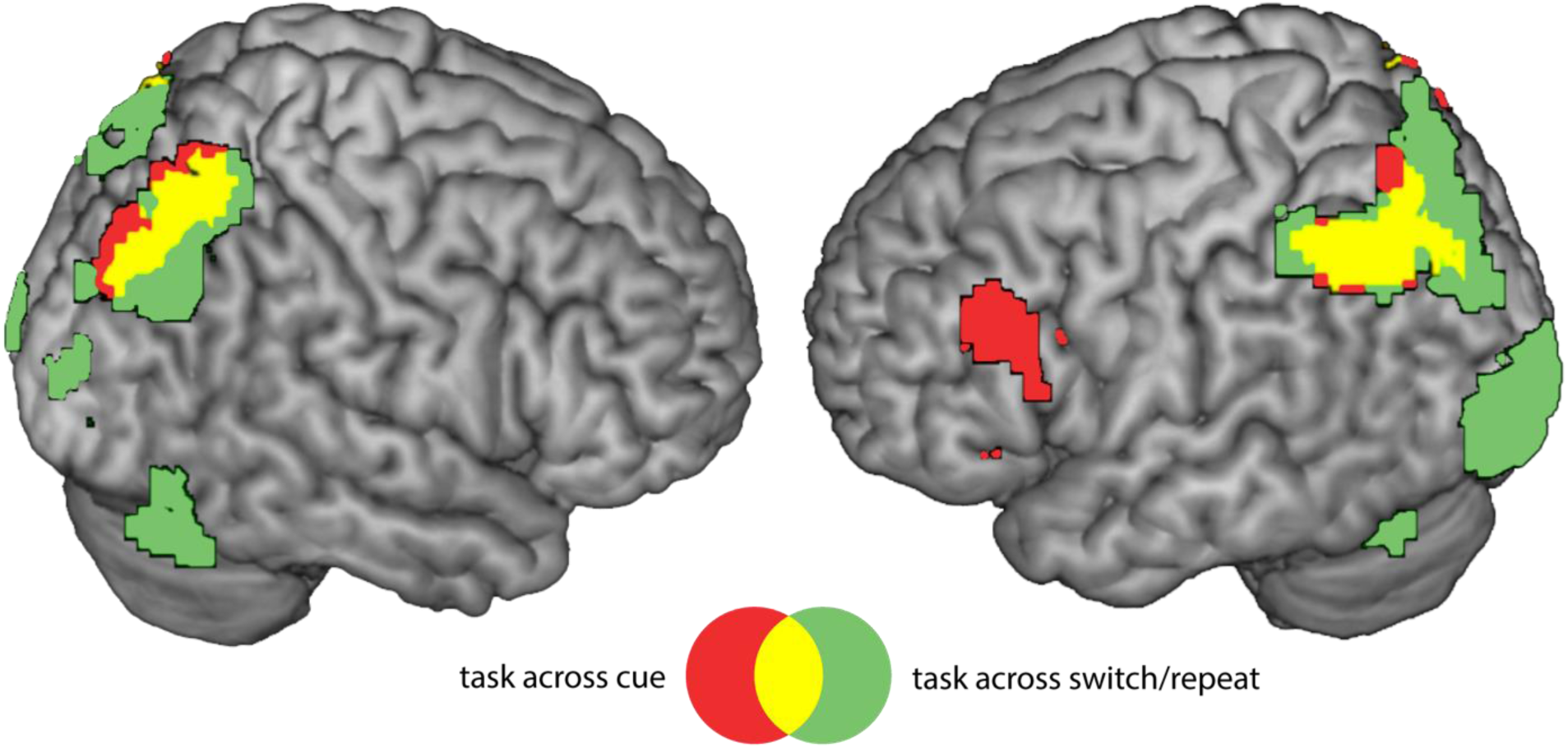
Task decoding in all subjects for the decoding of task across cues (red, analysis 1A) and the decoding of task across switch/repeat (green, analysis 2). Both regions overlap Note: task decoding across switch/repeat does not control for the visual information contained in task cues (which might explain occipital task information) and is also less sensitive than the leave-one-subject-ROI approach (which might account for no prefrontal cluster surviving cluster correction (for both analyses: *p* < 0.05, FWE corrected at the cluster level, voxel threshold *p* < 0.001).

## Discussion

### Summary

Effective goal directed behavior requires humans to frequently switch between different tasks. In order to direct this behavior, cognitive control is required (Kok et al., 2006). Much previous research used task switching paradigms to examine the role of cognitive control when changing between tasks (Kiesel et al., 2010; Monsell, 2003). Results show that performance is modulated by switching, and switch-costs are observed in both RT and accuracy (Allport et al., 1994; Jersild, 1927). Yet, the cognitive mechanisms and neuronal correlates of this behavioral switch cost are still under debate (Kiesel et al., 2010). Most previous fMRI research has focused on the neural correlates of task switching processes (Ruge et al., 2011) and task switching related processes have been associated with a fronto-parietal control network (Brass, Derrfuss, Forstmann, & von Cramon, 2005; Braver, Reynolds, & Donaldson, 2003; Crone, Wendelken, Donohue, & Bunge, 2006; Holmes et al., 2001). However, most of this research focused on the processes required to reconfigure the cognitive system from performing one task to performing a different task. Presumably, this includes changes to the neural representations of tasks, but effects on neural task representations have rarely been investigated before (but see Waskom et al., 2014). However, task representations have been shown to be context-dependent in some cases (Woolgar et al., 2015), while remaining context-independent in others (Wisniewski et al., 2016). Here, we investigated the influence of cognitive control processes related to task switching on the neural representations of tasks.

In the current study, subjects were cued to perform one of two simple tasks, with the task being repeated or switched between successive trials. Behavioral results indicate that subjects showed switch costs (Rogers & Monsell, 1995), which suggests cognitive control demands differed between switch and repeat trials. We first compared task decoding accuracies in switch and repeat trials in these regions. Our results show that tasks were represented in bilateral parietal cortex and left lateral PFC. However, we found no differences in task decoding accuracies between switch and repeat trials. Thus, our data yielded no evidence that tasks are represented differently for either switch or repeat trials in the regions that we previously identified to maintain task information (but see Waskom et al., 2014). We also tested for similarities and in task coding across switch and repeat trials using cross-classification. Results indicate that the fronto-parietal cortex represents tasks irrespective of the current cognitive control demands in task switching and suggests that tasks are coded in a robust, switching independent pattern.

### Task representations in fronto-parietal cortex

Recent MVPA research directly investigating the neural representations of tasks has shown that parietal (Bode & Haynes, 2009; Etzel, Cole, Zacks, Kay, & Braver, 2015; Waskom et al., 2014; Wisniewski, Reverberi, Momennejad, et al., 2015; Woolgar, Thompson, Bor, & Duncan, 2011) as well as medial PFC (Gilbert, 2011; Momennejad & Haynes, 2013) and lateral PFC (Cole et al., 2011; Reverberi et al., 2012b) hold information about tasks. We provide further evidence for these findings as we were able to discriminate between the two highly similar tasks in bilateral parietal and left lateral PFC. This is in line with previous results which highlight the important role of these regions in task processing during task retrieval and maintenance (Bunge, Kahn, Wallis, Miller, & Wagner, 2003; Gilbert, 2011; Sakai & Passingham, 2003), processing rule and task compositionality (Reverberi et al., 2012a; Woolgar, Thompson, et al., 2011), adaptively coding tasks under different conditions (Woolgar, Hampshire, et al., 2011) and their engagement over the course of development (Wendelken, Munakata, Baym, Souza, & Bunge, 2012).

### Influence of switching on task representation in fronto-parietal cortex

Recent studies suggest that task representations can be modulated by different contextual variables: task representations have been observed to be modulated by rule complexity (Woolgar et al., 2015), rewards (Etzel et al., 2015) or skill acquisition (Jimura et al., 2014). This illustrates how higher cognitive functions might flexibly change the way task are processed in the brain, possibly reflecting adaptation of neuronal populations to different environmental demands (Duncan, 2010, 2013). However, other studies suggest that task representations also remain unaffected by experimental manipulation, such as task novelty (Cole et al., 2011), task difficulty (Wisniewski, Reverberi, Tusche, & Haynes, 2015), or whether they are freely chosen or externally cued (Wisniewski et al., 2016; Zhang et al., 2013). It remains an open question whether and how cognitive control processes modulate task representations. In a previous study, Waskom et al. (2014) found task information in the inferior frontal, and intraparietal sulcus, as well as the occipito-temporal cortex. They found representations of rules regarding perceptual discriminations to be modulated by task switching, as they had the highest decoding accuracy after a task switch. Such effects on context information might be driven by attentional processes (Liu & Hou, 2013). Also note that Waskom et al. did not observe behavioral switch costs. It thus remains unclear whether cognitive control demands differed between switch and repeat trials, and whether these neuroimaging results in fact reflect control-related processes. In contrast, our subjects did show switch costs, indicating different control demands between switch and repeat trials. Importantly, as we presented task cues simultaneously with the task stimuli, participants could not prepare in advance for the new task on switch trials. Thus, switch costs presumably reflect both effects of task-set inertia and proactive interference, as well as increased control demands due to the requirement to retrieve and implement the new task-set and to reconfigure stimulus-response accordingly. Nevertheless, our results suggest that control demands do not modulate task representations. Taken together, these findings indicate that tasks are represented using a general, context-independent neural code. At first glance, this finding might be taken to imply that these brain regions do not support flexible adaptation of behavior, as they do not flexibly change under varying environmental conditions. It has previously been argued that frontal and parietal brain regions support flexible adaptation through flexible task representations which change under varying external demands (Duncan, 2001; Waskom et al., 2014; Woolgar et al., 2015). However, similar coding under different conditions might also support adaptive behavior: invariant coding allows robust access to task information even if we are confronted with novel situations. This might enable fast transfer of abstract rules (Cole et al., 2011) and stable selective attention towards task-relevant information (Zhang et al., 2013). Stable task representations have also been observed under varying attentional loads (Chan, Kucyi, & DeSouza, 2015), further highlighting the context-independent coding of tasks. Thus, our findings of such invariant neural representations do not rule out a dynamic adjustment of task specific neurons, as the adaptive coding hypothesis (Duncan, 2001, 2010; Waskom et al., 2014) suggests. Flexible top-down signals may be reflected in different levels of task processing that merely access the robust context-independent representation without modulating it. Additionally, we found no significant results in the analysis testing for context dependent task coding. Although we used a highly sensitive ROI approach, this null finding cannot rule out in principle that there might also be neurons that do code tasks differently for different cognitive control demands.

### Role of task switch processes

Although this study focused on differences and similarities of neural task representations during switching, we also observed behavioral switch costs. Our paradigm was not designed to determine the source of the underlying processes, but switch costs might arise for a number of reasons, including proactive interference due to task-set inertia (Allport et al., 1994), the inhibition of previously executed task-sets (Goschke, 2000; Mayr & Keele, 2000), and processes of rule retrieval (*goal setting*) and rule implementation (Rubinstein, Meyer, & Evans, 2001). Models of task switching which assume that part of the switch cost reflects proactive interference from previous and/or crosstalk from concurrently active, but currently irrelevant task-sets, would presumably result in task representations that are degraded and less distinct on switch compared to repeat trials. Such an effect should show up in a reduced accuracy with which task representation can be decoded from spatial patterns of brain activity. However, the present findings of task representations that are independent of current switch demands do not suggest such a modulation, from whichever source. Neurons in the fronto-parietal cortex are able to encode tasks similarly under various different conditions, like high and low control demands (see also Wisniewski et al., 2016). Switch costs might then arise at a different stage, when task information from the parietal cortex is further processed by brain regions more closely associated with implementing cognitive control (Badre, 2008).

### Conclusion

In summary, our results provide novel insights into the effects of task switching on the distributed neuronal representations of tasks. We did not find reliable differences in task coding between switch and repeat trials. However, task representations in bilateral parietal and left prefrontal cortex were similar under conditions of high and low cognitive control demands. These results provide further insight into the important function of the fronto-parietal network for task representation. Control-independent task coding might enable robust access to task-relevant information under different environmental conditions, in order to support flexible adjustment of behavior.

## Acknowledgements

We would like to thank Kai Görgen, Achim Meier, Jelle Demanet and Marcel Brass for their helpful comments on this project. This work was supported by the Bernstein Computational Neuroscience Program of the German Federal Ministry of Education and Research (grant reference 01GQ1001C) and the German Research Foundation within the Collaborative Research Center “Volition and Cognitive Control: Mechanisms, Modulations, Dysfunctions” (DFG grants SFB 940/1 and SFB 940/2). It was further supported by German Research Foundation grants Exc 257, Neurocure and KFO247, and Research Foundation Flanders grant FWO.OPR.2013.0136.01.

